# High-throughput developmental assay of cold tolerance in *Caenorhabditis elegans*

**DOI:** 10.1101/2025.09.14.676146

**Authors:** Amanda L. Peake, Nikita S. Jhaveri, Erik C. Andersen, John R. Stinchcombe

**Affiliations:** Department of Ecology and Evolutionary Biology, University of Toronto, 25 Willcocks St, Toronto, Ontario, Canada, M5S3B2; Department of Biology, Johns Hopkins University, Baltimore, Maryland, United States

## Abstract

Temperature can impose strong selection causing thermal tolerance variation between individuals, populations, and species. We developed a high-throughput larval development assay for cold tolerance in the model organism *Caenorhabditis elegans*. We exposed animals to 4°C cold treatments for either 12 or 24 hours. Animals exposed to the 24-hour cold treatment exhibited greater variation and heritability in cold tolerance during the L1 larval stage. The high-throughput approach that we developed is easily scalable to simultaneously measure a large number of strains, which makes it ideal for studying the genetics and evolution of cold tolerance in *Caenorhabditis* nematodes.

## Description

Phenotypic variation within and between populations can provide insights into the genetic basis and evolutionary history of ecologically important traits. Broadly distributed species must adapt to local environmental conditions to avoid extinction. In turn, populations are expected to harbor beneficial genetic variants that contribute to locally adapted phenotypes (Hedrick 2006). Temperature can be a strong selective pressure and temperature differences across a species range can have major evolutionary consequences (Huey and Kingsolver 1989; Williams et al., 2015; Sunday et al., 2019).

We developed a high-throughput developmental assay to study cold tolerance in *Caenorhabditis elegans*. The species occupies a wide geographic range, and wild isolates have been sampled at substrate temperatures ranging from 3.9°C to 26°C (Cook et al., 2017; Crombie et al., 2019, 2022, 2024). With an ever increasing number of wild isolates being sampled and sequenced by the *Caenorhabditis* Natural Diversity Resource (CaeNDR), it is now possible to study naturally occurring variation (Cook et al., 2017; Crombie et al., 2024). Accurately characterizing phenotypic and genetic variation requires large sample sizes evaluated in a common environment. Therefore, high-throughput phenotyping methods are required to quantify variation in dozens to hundreds of strains.

As a proof of concept, we developed a high-throughput phenotypic assay for cold tolerance and evaluated it using seven *C. elegans* strains (CB4856, CX11314, DL238, ECA1286, JU394, JU1200, and N2). We exposed animals at an early developmental stage (L1s) to 4°C for either 12 or 24 hours. Previous cold tolerance studies in *Caenorhabditis* nematodes have mostly focused on adults (Robinson and Powell 2016; Wang et al., 2021; Vigne and Braendle 2025), however, thermal tolerance can differ between adult and larval stages (Jiang et al., 2018; Jhaveri et al., 2025a). We used 12 and 24 hours to mimic daily temperature fluctuations in nature. The 4°C cold treatment is ecologically relevant because 3.9°C is the lowest sampling substrate temperature of CaeNDR wild strains (CaeNDR release 20250626; Cook et al., 2017; Crombie et al., 2024). After the cold treatment, animals were fed and returned to 20°C for 48 hours. Animals increase in length throughout development, therefore, we used animal length as a proxy for developmental rate. We quantified cold tolerance as differences in length between cold-treated animals and control animals kept at 20°C. Strains with larger differences in length between cold-treated and control animals indicate lower cold tolerance.

In the 12-hour cold treatment, cold-treated individuals had a reduced developmental rate but the cold treatment impacted most strains similarly (Figure 1A). Cold-treated animals were significantly smaller than the control (two-way type III ANOVA: F_treatment_(1,6) = 98.98, *p*_treatment_ < 0.001) and differences among strains explained a significant amount of variation in animal length (likelihood ratio test: χ^2^_strain_ (1) = 3.94, *p*_strain_ = 0.02). Although differences in how different strains responded to cold treatment also explained a significant amount of variation in animal length (likelihood ratio test: χ^2^_strain_ (1) = 89.22, *p*_strain:treatment_ < 0.001), we observed mostly parallel reaction norm slopes except for the strain DL238 (Figure 1A). Strains had a mean difference of 76 - 151 µm between cold-treated and control animals (Figure 1A). The broad-sense heritability for animal length across both control and treatment individuals (*H*^2^_strain_) was 0.47 and the broad-sense heritability in how animal length changes in response to cold treatment (*H*^2^_strain:treatment_) was 0.2 (Figure 1A). Therefore, the 12-hour cold treatment had a similar effect on developmental rate for most strains so we did not observe high heritability in cold tolerance among strains for the 12-hour cold treatment.

**Figure 1.**
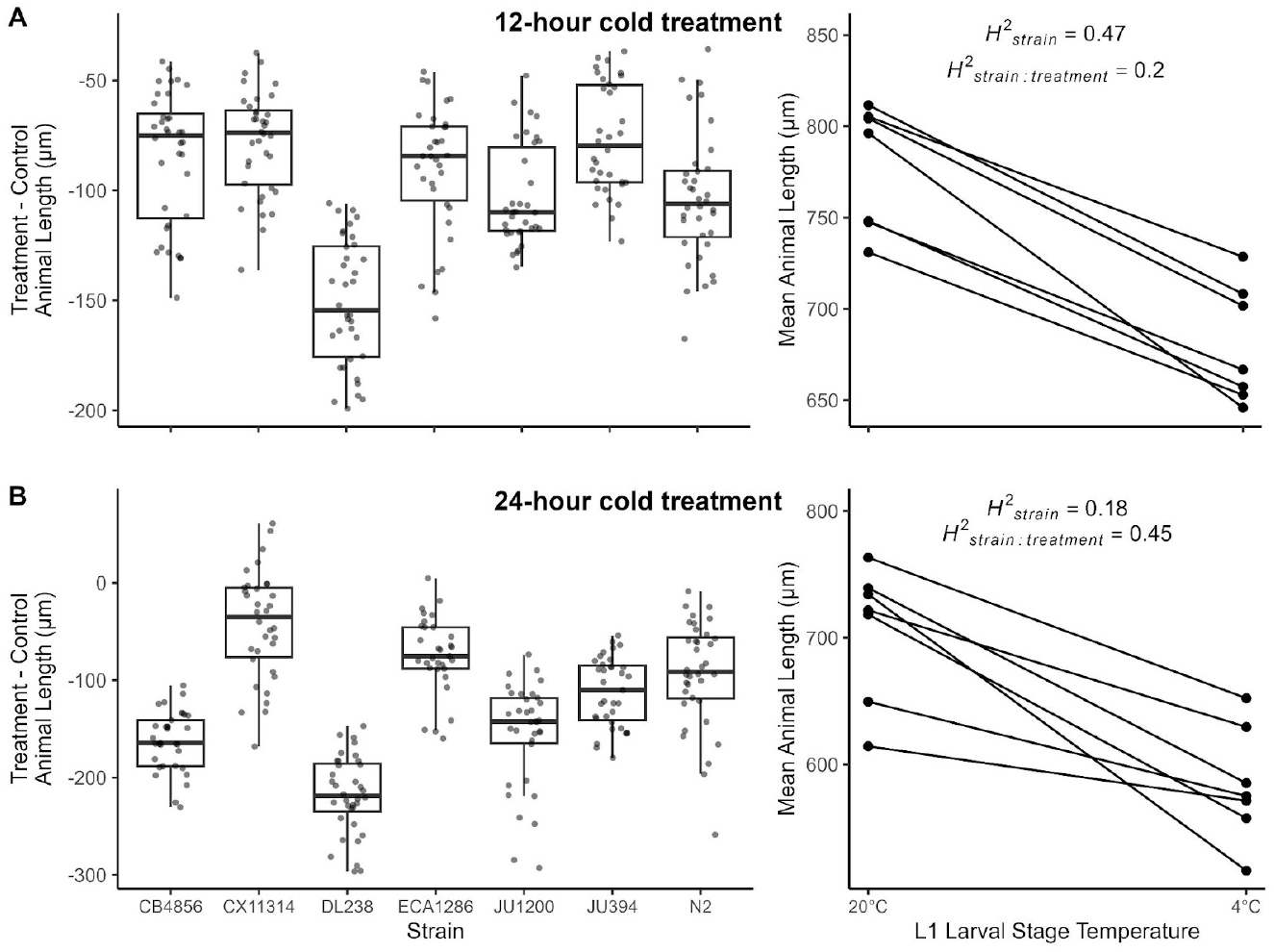
Effect of 4°C cold treatment on development of seven *C. elegans* strains: Cold treatment effects on development for the duration **A)** of 12 hours or **B)** of 24 hours. Box plots of difference in animal length (µm) between 4°C cold-treated animals and 20°C control animals are shown on the left. A larger difference in length between cold-treated animals and the control indicates a more severe effect of cold treatment on development. Reaction norm plots for animal length (µm) from control treatment (20°C) and cold-treated (4°C) animals are shown on the right. Data points in the reaction norm plots represent strain mean values. *H*^2^_strain_ is the broad-sense heritability for animal length across both environments. *H*^2^_strain:treatment_ is the broad-sense heritability for the plasticity in length, or how much length changes in response to cold treatment.

In the 24-hour cold treatment, animal length significantly differed between the control and cold treatment (two-way type III ANOVA: F_treatment_(1,6.01) = 29.25, *p*_treatment_ = 0.002). Although differences among strains did not explain a significant amount of variation in animal length (likelihood ratio test; χ^2^_strain_(1) = 0.48, *p*_strain_ = 0.25), we found a significant amount of variation was explained by strains responding differently to the cold treatment (likelihood ratio test: χ^2^ _strain:treatment_ (1) = 180.45, *p*_strain:treatment_ < 0.001). Mean differences between cold-treated and control animals ranged from 43 µm to 218 µm and reaction norm slopes were mostly non-parallel among strains (Figure 1B). In the 24-hour cold treatment, *H*^2^_strain_ = 0.18 and *H*^2^_strain:treatment_ = 0.45 (Figure 1B), indicating that cold treatment impacted the developmental rate of strains differently and was heritable.

We found heritable variation in how developmental rate was impacted by cold exposure at early larval stages in *C. elegans*. Out of the two cold treatment durations, we recommend the 24-hour duration because it had a higher heritability of how developmental rate is affected by cold treatment (*H*^2^_strain:treatment_). The high-throughput assay can be easily scaled to include a large number of strains, and therefore, provides an avenue for exploring the genetics and the evolution of cold tolerance in *C. elegans* and other closely related species (Widmayer et al., 2022b).

## Methods

We used seven *C. elegans* strains (N2, ECA1286, DL238, CB4856, CX11314, JU394, and JU1200) from CaeNDR (Cook et al., 2017; Crombie et al., 2024). Strains were thawed and maintained for three generations to reduce any transgenerational effects of starvation. We maintained strains at 20°C on 6 cm NGMA plates with 1% agar and 0.7% agarose (Andersen et al., 2014) seeded with *Escherichia coli* strain OP50. We age synchronized animals using filtration (Jhaveri et al., 2025b) and suspended embryos in K medium to a final embryo concentration of 1 embryo/µL. We distributed 50 µL embryo solution into wells in 96-well plates, where each strain had one row of 12 wells with approximately 50 animals per well. Cold treatments and controls had three replicate plates with three different plate designs to account for edge effects. We then placed plates in a humidity chamber within a 20°C shaking incubator to get a synchronized L1 larval population. Shaking incubators throughout the experiment were set to 170 rpm.

To determine the effects of cold temperature on development, we exposed L1 animals to 4°C for either 12 hours or 24 hours. Treatment plates were placed in a shaking incubator set to 4°C and control plates remained in a shaking incubator set to 20°C. After the cold treatment, we fed animals 25 µL of OD_600_30 *E. coli* strain HB101. We prepared food from thawed aliquots of OD_600_100 HB101 diluted with K medium and 150 µM kanamycin to avoid contamination (Widmayer et al., 2022a). After feeding, we placed plates in the 20°C shaking incubator for 48 hours. To straighten animals for automated image analysis, we paralyzed animals with 334 µL of 50 mM sodium azide and imaged plates 10 minutes later using a Molecular Devices ImageXpress Nano microscope with a 2x objective lens (Shaver et al., 2023).

We processed images using CellProfiler (v4.2.8) to extract animal lengths (Carpenter et al., 2006; Wählby et al., 2012; Widmayer et al., 2022a). The CellProfiler pipeline categorises objects as L1-L4 animals and also includes a mulit-drug high dose (MDHD) category that identifies extremely small animals. We used easyXpress (v2.0.0) in R (v4.4.0) to clean the processed image data (Nyaanga et al., 2021; R Core Team 2024). Upon visual inspection of the wells, none of the wells had extremely small animals that should fall into the MDHD category. We excluded MDHD objects and objects < 165 µm that are likely debris incorrectly detected as animals. We also excluded wells that had fewer than five or more than 60 animals. We then calculated median animal lengths per well and excluded outlier wells where median lengths fell outside +/-1.5 times the interquartile range of wells grouped by strain, treatment/control, and replicate plate. To obtain a proxy of cold tolerance, we subtracted the median animal length for each treatment well from the mean of the medians of the control wells for individual strains for each plate design.

We used a two-way type III ANOVA with Satterthwaite’s method of approximation using the model below for each cold treatment to determine if animal length differed between the cold treatment and the control. We used a likelihood ratio test on the same model to determine if the strain and strain:treatment terms were significant and halved the p-values to correct for testing random effects (Kuznetsova et al., 2015). Statistical tests were done using the lmerTest R package (v. 3.1-3; Kuznetsova et al., 2017).

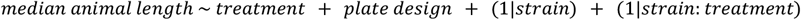

We calculated broad-sense heritability for animal length across environments (*H*^2^_strain_) and the broad-sense heritability for how animal length changes in response to the cold treatment (*H*^2^_strain:treatment_) by extracting variance components from the linear model above (Scheiner and Lyman 1989).

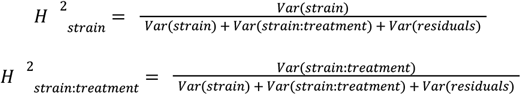

We visualized data in R using ggplot2 (v3.5.2; Wickham 2016), gridExtra (v2.3; Auguie 2017), cowplot (v1.1.3; Wilke 2025) and ggpubr (v0.6.0; Kassambara 2023). Scripts available: https://github.com/peakeama/C.elegans_cold_tolerance

## Supporting information

Animal length data from images processed using CellProfiler and easyXpress for 12 hour cold treatment

Animal length data from images processed using CellProfiler and easyXpress for 24 hour cold treatment

## Author contributions

Amanda L. Peake: conceptualization, data curation, formal analysis, funding acquisition, investigation, methodology, visualisation, validation, writing - original draft

Nikita S. Jhaveri: data curation, formal analysis, investigation, methodology, writing - review & editing

Erik C. Andersen: resources, funding acquisition, supervision, writing - review & editing John R. Stinchcombe: funding acquisition, supervision, writing - review & editing

## Acknowledgements

We would like to thank members of the Andersen Laboratory for feedback on the project and Dr. Asher Cutter (Univ. of Toronto) for facilitating the collaboration.

## Funding

This work was supported by a University of Toronto Center of Global Change Science Graduate Student Award to Amanda L. Peake. We would also like to thank NSERC for the funding provided as part of a PGS D award to Amanda L. Peake and a Discovery Grant to John R. Stinchcombe. This work was also supported by an NSF CAREER award to Erik C. Andersen (1751035).

## Notes

https://github.com/peakeama/C.elegans_cold_tolerance

### Competing Interest Statement

The authors have declared no competing interest.

